# Molecular evidence of intertidal habitats selecting for repeated ice-binding protein evolution in invertebrates

**DOI:** 10.1101/2021.08.30.458284

**Authors:** Isaiah C. H. Box, Benjamin J. Matthews, Katie E. Marshall

## Abstract

Ice-binding proteins (IBPs) have evolved independently in multiple taxonomic groups to improve their survival of sub-zero temperatures. Intertidal invertebrates in temperate and polar regions frequently encounter sub-zero temperatures, yet there is little information on IBPs in these organisms. We hypothesized that there are far more ice-binding proteins than are currently known and that the occurrence of freezing in the intertidal zone selects for these proteins. We compiled a list of genome-sequenced invertebrates across multiple habitats and a list of known IBP sequences and used BLAST to identify a wide array of putative IBPs in those invertebrates. We found that the probability of an invertebrate species having an ice-binding protein was significantly greater in intertidal species as compared to those primarily found in open ocean or freshwater habitats. These intertidal IBPs had high sequence similarity to fish and tick antifreeze glycoproteins and fish type II antifreeze proteins. Previously established classifiers based on machine learning techniques further predicted ice-binding activity in the majority of our newly identified putative IBPs. We investigated the potential evolutionary origin of one putative IBP from the hard-shelled mussel *Mytilus coruscus* and suggest that it arose through gene duplication and neofunctionalization. We show that IBPs likely readily evolve in response to freezing risk, that there is an array of uncharacterized ice binding proteins and highlight the need for broader laboratory-based surveys of the diversity of ice binding activity across diverse taxonomic and ecological groups.

**Summary statement:** Intertidal invertebrates have a disproportionate number of putative ice-binding proteins relative to other habitats. These putative proteins are highly similar to antifreeze glycoproteins and type II antifreeze proteins from fish.

## INTRODUCTION

For organisms to inhabit temperate and polar regions, low temperature tolerance is a key trait (Sanmartín et al., 2001; Wiens and Donoghue, 2004). This is especially true for ectotherms that risk freezing of their body fluids and therefore have evolved strategies to survive otherwise lethal sub-zero body temperatures (Lee, 2010). While the physiology and biochemistry of low temperature tolerance is diverse, one common biochemical mechanism used by a broad array of organisms ranging from bacteria to fish are ice-binding proteins (IBPs; Davies, 2014; Bar Dolev et al., 2016).

As the name suggests, IBPs are proteins that can bind to ice (Davies, 2014; Bar Dolev et al., 2016). How these proteins bind to ice is poorly understood, but one hypothesis is that ice binding is accomplished through hydrogen bonding and Van der Waals forces (Jia et al., 1996; Bar Dolev et al., 2016). Regardless of the mechanisms, these proteins promote survival at sub-zero temperatures through three primary means. First, ice binding can prevent further ice formation by effectively suppressing the freezing point relative to the melting point, resulting in thermal hysteresis (Davies, 2014; Bar Dolev et al., 2016). Thermal hysteresis is advantageous for freeze-avoidant organisms but can also be advantageous for freeze tolerant organisms that want to prevent ice formation in particularly sensitive tissues or intracellular space (Davies, 2014; Bar Dolev et al., 2016). Ice binding can also prevent changes in the size of ice crystals through ice recrystallization inhibition (IRI; Davies, 2014; Bar Dolev et al., 2016). Ice recrystallization is the thermodynamically driven process whereby many small ice crystals present in early ice matrices incorporate into each other to form fewer but larger ice crystals (Pronk et al., 2005; Balcerzak et al., 2014). This growth of ice crystals in the body can damage cells and tissues, thus the IRI activity of IBPs is important for surviving internal ice formation (Davies, 2014; Bar Dolev et al., 2016). Finally, IBPs can nucleate ice, promoting the formation of ice in the body (Davies, 2014; Bar Dolev et al., 2016). Ice nucleation activity can allow organisms to control when and where ice forms in the body, preventing uncontrolled ice formation in the body which could increase the risk of lethal freezing injury (Davies, 2014; Bar Dolev et al., 2016).

The variety of ways IBPs influence ice formation is mirrored by the diversity of organisms that utilize them, with examples of IBPs found throughout the tree of life (Bar Dolev et al., 2016). For example, IBPs with ice nucleation activity have been described in bacteria (Xu et al., 1998; Ling et al., 2018), with thermal hysteresis activity in fish (Fletcher et al., 2001), and with IRI activity in plants (John et al., 2009; reviewed in Bar Dolev et al., 2016). In fact, this list is ever-expanding, and novel IBPs are regularly discovered (Scholl et al., 2021). IBPs have evolved independently in multiple taxonomic groups resulting in distinct groups of IBPs from beetles (Coleoptera) and moths (Lepidoptera) and have even evolved independently multiple times within a single taxon, including three unique IBP lineages in rye grasses and five lineages in teleost fishes (Bildanova et al., 2012). This structural diversity coincides with an array of evolutionary pathways that can produce IBPs (Bildanova et al., 2012). In teleosts alone there are examples of IBPs evolving *de novo* (Zhuang et al., 2019), through horizontal gene transfer (Graham et al., 2008), and through neofunctionalization (Deng et al., 2010). There are multiple progenitor proteins that have evolved into IBPs including C-type lectins for type II antifreeze proteins (AFPs) of fish (Gronwald et al., 1998), trypsinogen-like proteases for fish antifreeze glycoproteins (AFGPs; Chen et al., 1997), and multiple others (summarized in Bildanova et al., 2012). This combined diversity of functions, structures, evolutionary origins, and organisms of origin for IBPs point to a relative ease of evolving an IBP for surviving temperate and polar habitats.

Despite this ready evolution of IBPs across the tree of life, there is a vast polyphyletic group that is thus far largely devoid of suspected IBPs: intertidal invertebrates (Storey and Storey, 2013). This is surprising as intertidal invertebrates in temperate regions experience freezing temperatures in the winter during low tide, and many species, especially the slow-moving and sessile ones, are freeze tolerant (Aarset, 1982). While it is possible for an organism to tolerate freezing without the use of IBPs, other known molecular strategies of freeze tolerance such as accumulating polyols and sugars are not used by intertidal invertebrates (Storey and Storey, 2013). While osmolyte accumulation is an important correlate to freeze tolerance in the bay mussel (*Mytilus trossulus*), an intertidal species, it does not fully explain its freeze tolerance (Kennedy et al., 2020). This suggests high molecular weight cryoprotectants such as IBPs may play a role in their freeze tolerance (Kennedy et al., 2020). In addition to this, there has been evidence for IBPs in both blue mussels (*Mytilus edulis*) and barnacles (*Semibalanus balanoides*; Theede et al., 1976; Marshall et al., 2018 preprint) and a partially characterized IBP with ice nucleating activity found in an intertidal air-breathing snail (Madison et al., 1991). This, combined with the current scarcity of research on freeze tolerance in intertidal invertebrates suggests that there is an array of IBPs waiting to be discovered in the intertidal zone (Storey and Storey, 2013), especially given the stressful habitat they reside in and the relative ease of evolving IBPs.

We hypothesized that IBPs are widespread and unreported in intertidal invertebrates, and that the intertidal habitat has selected for the evolution of IBPs. Due to examples of highly similar IBPs being acquired both convergently (Zhuang et al., 2019) and through horizontal gene transfer (Graham et al., 2008), we predicted that a BLAST search using sequenced and laboratory-characterized IBPs would return strong matches in the molecular sequence data of intertidal invertebrates. Expanding on this, we predicted that IBP matches would be biased towards species that occur in the intertidal zone relative to species in the same phylum found in different habitats. As a result of these investigations, we systematically demonstrate that putative IBPs are broadly distributed through invertebrate taxa, that they were identified through sequence homology to type II AFPs, AFGPs, and Coleopteran AFPs, and that intertidal species are more likely to contain these putative IBPs than their non-intertidal conphyletics. Taken together, this suggests that IBPs readily evolve along the same evolutionary trajectory in response to low temperature stress.

## METHODS

### Data Collection

To investigate the presence of putative IBPs in intertidal invertebrates, we first compiled a query list of known IBPs. Query list IBPs met two criteria: First, the IBP had to have amino acid sequence information available with an NCBI accession number. Second, the IBP had to have literature documentation of ice-binding activity (Table S1).

Following the creation of this IBP query list, we produced a list of organisms to search for putative novel IBPs. We selected the following phyla of common intertidal invertebrates: Mollusca, Crustacea (subphylum of Arthropoda), Echinodermata, Annelida, and Cnidaria. We opted only for the subphylum Crustacea since the vast majority of genome-sequenced arthropods are terrestrial. We then compiled a list of all species within these (sub)phyla that had a sequenced genome available at NCBI (NCBI, 1988), and classified the habitat type of each search organism into “terrestrial”, “endoparasitic”, “intertidal”, “subtidal”, “estuarine”, or “other marine” based on listings on SeaLifeBase (Palomares and Pauly [editors], 2020), WoRMS (Horton et al., 2021), and direct citations (Table 1; Table S2). In this study, subtidal refers only to the shallow subtidal (≤ 1 m; Saier, 2002) while “other marine” is a blanket term used in this study for all ocean habitats that do not fit within the intertidal, shallow subtidal, or estuarine habitat types. For species found in a wide range of ocean habitats, their shallowest habitat type was used for categorization in this study.

**Table 1.**
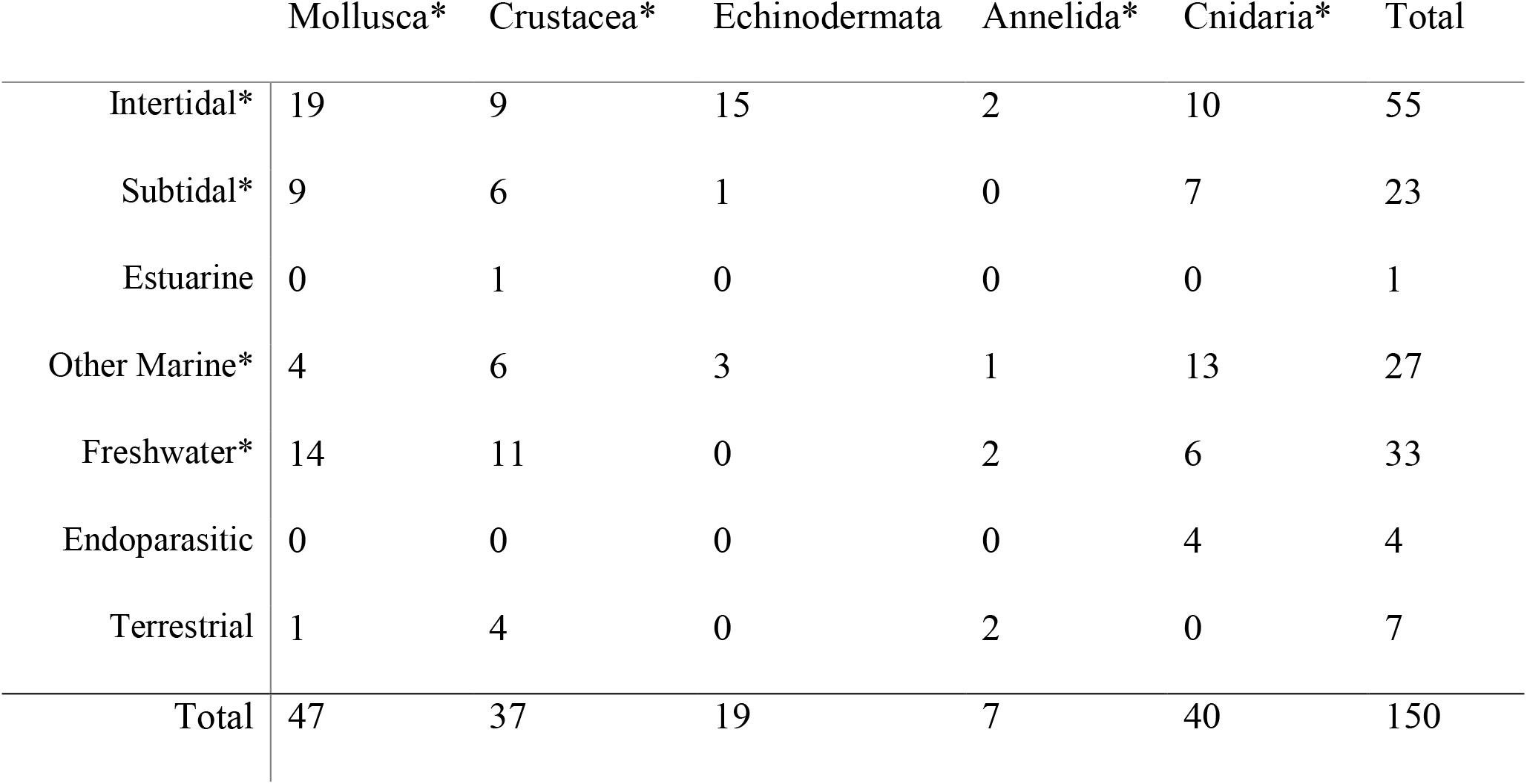
Distribution of search organisms by (sub)phylum and habitat type. Asterisk indicates predictors used in logistic regression examining predictors of IBP presence.

### Search for Putative IBPs

As there are no common ice-binding domains across all IBP types (Davies, 2014; Bar Dolev et al., 2016), we used the protein BLAST algorithm to identify any sequence homology between the query IBP list and the organism search list (Altschul et al., 1990). A protein BLAST (blastp) search was performed on the non-redundant protein sequence database filtered only to include the species in our organism search list. We set the expect threshold to an E-value of 1 × 10^−5^ with a word size of six amino acids and enabling automatic adjustments for short query sequences. We used the standard BLOSUM62 matrix with conditional compositional score matrix adjustment method, gap existence cost of 11, and extension set to 1 to determine alignment scores (Henikoff and Henikoff, 1992). We binarily categorized each organism in the search list based on if they contained an IBP hit conforming to the 1 × 10^−5^ expect threshold from this search for later comparison of IBP presence among habitats and phyla.

To visualize potential phylogenetic patterns in IBP hits, we produced an unrooted phylogeny from the IBP query sequences. Query sequences were aligned in MEGA (Kumar et al., 2018) using the MUSCLE algorithm (Edgar, 2004). A maximum likelihood tree was produced from the aligned sequences using the WAG+G+F model (Whelan and Goldman, 2001). Our model selection for this and all future phylogenies in this study was determined by calculating maximum-likelihood fits in MEGA (Kumar et al., 2018) for 56 different amino-acid substitution models, then selecting the model with the lowest Bayesian Information Criterion. We then mapped search organisms containing BLAST matches with high homology to query list IBPs on the tree based on habitat and phylum to guide our investigations into the potential evolutionary origin of the putative IBPs. We also produced a phylogeny of the organism search list through the taxonomy browser in NCBI, highlighting species bearing BLAST matches for IBPs to gain further insight into the evolution of putative IBPs.

### Assessing Putative IBPs

We then investigated the clades of query list IBPs that contained multiple hits against the organism search list to ensure matches were not due to ubiquitous IBP-like sequences found across organisms, including the progenitor proteins of the IBPs. To do this, sequences in each IBP clade were realigned using CLUSTAL W (Thompson et al., 1994) as in Arning (2018). We produced new maximum likelihood trees from these alignments, each using 50 bootstrap replicates and the WAG+G, WAG+G+F (Whelan and Goldman, 2001) and JTT (Jones et al., 1992) models for the Coleopteran AFP, AFGP, and Teleost type II AFP clades, respectively. In the stead of an ice-binding domain search, we calculated ancestral sequences from these trees through MEGA (Kumar et al., 2018) using the same respective models mentioned above to obtain protein sequences bearing sequence regions shared across the respective IBPs in the clade. We searched for these ancestral sequences across NCBI (1988) using blastp on the non-redundant protein sequence database to investigate whether these had higher homology for IBPs and not the progenitor proteins of IBPs (Altschul et al., 1990). In the case of arthropod IBPs, progenitor proteins are unknown (Bildanova et al., 2012) but since the tick AFGPs are similar in sequence to fish AFGPs (Neelakanta et al., 2010), we assumed the same trypsinogen-like protease progenitor protein (Chen et al., 1997). We then repeated the protein BLAST on the organism search list using the ancestral sequences to confirm a similar match profile with respect to which organisms bear matches (Altschul et al., 1990). We compiled hits from the ancestral sequence to the organism search list and used them to obtain separate trees and ancestral sequences as above to repeat the above progenitor protein verification. We searched for these ancestral sequences from the search organisms across NCBI’s (1988) non-redundant protein sequence database using blastp to ensure a lack of matches for IBP progenitors or ubiquitous sequences.

To further assess the identified putative IBPs, we used a series of published machine learning algorithms for classifying IBPs. All matches were assessed this way except in the case of the type II AFPs and the carrot (*Daucus carota*) AFP which instead had the match with the lowest e-value for each search organism assessed. This was because these query IBPs had too many matches to assess given the limitations on the calculation time for some of the calculators used. The three web-accessible calculators we used for predicting AFPs through machine learning were: TargetFreeze (He et al., 2015), iAFP-Ense (Xiao et al., 2016), and CryoProtect (Pratiwi et al., 2017). For unknown reasons, not all sequences yielded outputs from the IBP calculators; 2.0% of sequences had no outputs from TargetFreeze (He et al., 2015) and 4.0% from CryoProtect (Pratiwi et al., 2017). Type II AFPs were further tested using the web-based disulfide bond predictor DiANNA 1.1 (Ferrè and Clote, 2005a,b, 2006), because type II AFPs have five disulfide bridges while C-type lectins have fewer (Graham et al., 2008). For all calculators, we also input the query IBPs to determine how much confidence we can have in each calculator’s output.

### Investigating Evolutionary Origin of a Putative IBP

To gain some insight into how these potential intertidal IBPs evolved, we evaluated the presence of genomic synteny for the best putative intertidal IBP for the type II AFP clade that met the criteria for all the above calculators: an unnamed protein product from *Mytilus coruscus* (NCBI Accession: CAC5422424.1). We located the coding region for the putative IBP in *M. coruscus*’s genome noting the contig the putative IBP gene is in and the 4 nearest protein coding genes (Table S3). We then performed tblastn searches (default parameters) for our putative IBP on the genomes of species of different degrees of relatedness to *M. coruscus* to locate orthologs (Table 2; Altschul et al., 1990). The contigs bearing these orthologs were then noted for each of these species as above (Table S3). We then performed tblastn searches on each genome for the four proteins surrounding the coding region for our primary protein of interest, comparing the arrangements of the coding regions for these proteins relative to *M. coruscus* to evaluate presence of synteny. Genome snapshots (130,000 nt) surrounding the coding site for the potential orthologs of our putative IBP for each of the above species were then input into SimpleSynteny (Veltri et al., 2016) along with the sequences of our protein of interest and its four surrounding proteins to visualize gene arrangement among species. To better understand trends in the gene mapping data, our protein of interest and its surrounding unknown protein products were aligned in MEGA (Kumar et al., 2018) using the MUSCLE algorithm (Edgar, 2004). A maximum likelihood tree was produced from the aligned sequences using 50 bootstrap replicates under the WAG model (Whelan and Goldman, 2001).

**Table 2.** Species used in the evaluation of genomic synteny for a putative IBP in *Mytilus coruscus* and their relatedness to *M. coruscus*.

| Species | Shared Taxonomic<br>Level with *M.*<br>*coruscus* | Shared Taxon Name | Genome NCBI<br>Accession |
| --- | --- | --- | --- |
| *Mytilus coruscus* | Species | *Mytilus coruscus* | GCA_011752425.2 |
| *Mytilus galloprovincialis* | Genus | *Mytilus* | GCA_900618805.1 |
| *Perna viridis* | Family | Mytilidae | GCA_018327765.1 |
| *Limnoperna fortunei* | Family | Mytilidae | GCA_003130415.1 |
| *Pecten maximus* | Subclass | Pteriomorpha | GCA_902652985.1 |
| *Crassostrea gigas* | Subclass | Pteriomorpha | GCA_902806645.1 |
| *Sinonovacula constricta* | Class | Bivalvia | GCA_007844125.1 |
| *Elysia chlorotica* | Phylum | Mollusca | GCA_003991915.1 |

### Statistical Analysis

To determine if the presence of IBP BLAST hits in search organisms is determined by habitat, phylum, or the interaction between them, we used a logistic regression model. The sample sizes for terrestrial, estuarine and endoparasitic organisms were too small (n = 7, 1, and 4, respectively) to include in this statistical analysis and we therefore omitted them. We also had to omit Annelida due to small sample size (n = 7) and Echinodermata because they are not found in freshwater habitats which would impede the comparison across habitat types. To determine significant differences between groups we used Tukey’s honest significance test on the logistic regression model (Tukey, 1949). Statistical analysis was completed using R (R Core Team, 2019) and the multcomp package (Hothorn et al., 2008), and alpha was set to 0.05.

## RESULTS

### Data Collection and Search for Putative IBPs

The organism search list contained 150 species across the 5 (sub)phyla (Table 1; Table S2). Not all (sub)phyla were represented in all habitat types (Table 1). A total of 148 query IBP sequences were compiled with broad taxonomic representation reflecting the currently known diversity of IBP-producing taxa (Bar Dolev et al., 2016; Fig. 1; Table S1). Using blastp, we found that 42 of the query sequences had high homology with at least one protein sequence in 53 of the species in the search list (Fig. 1; Fig. 2).

**Fig. 1.**
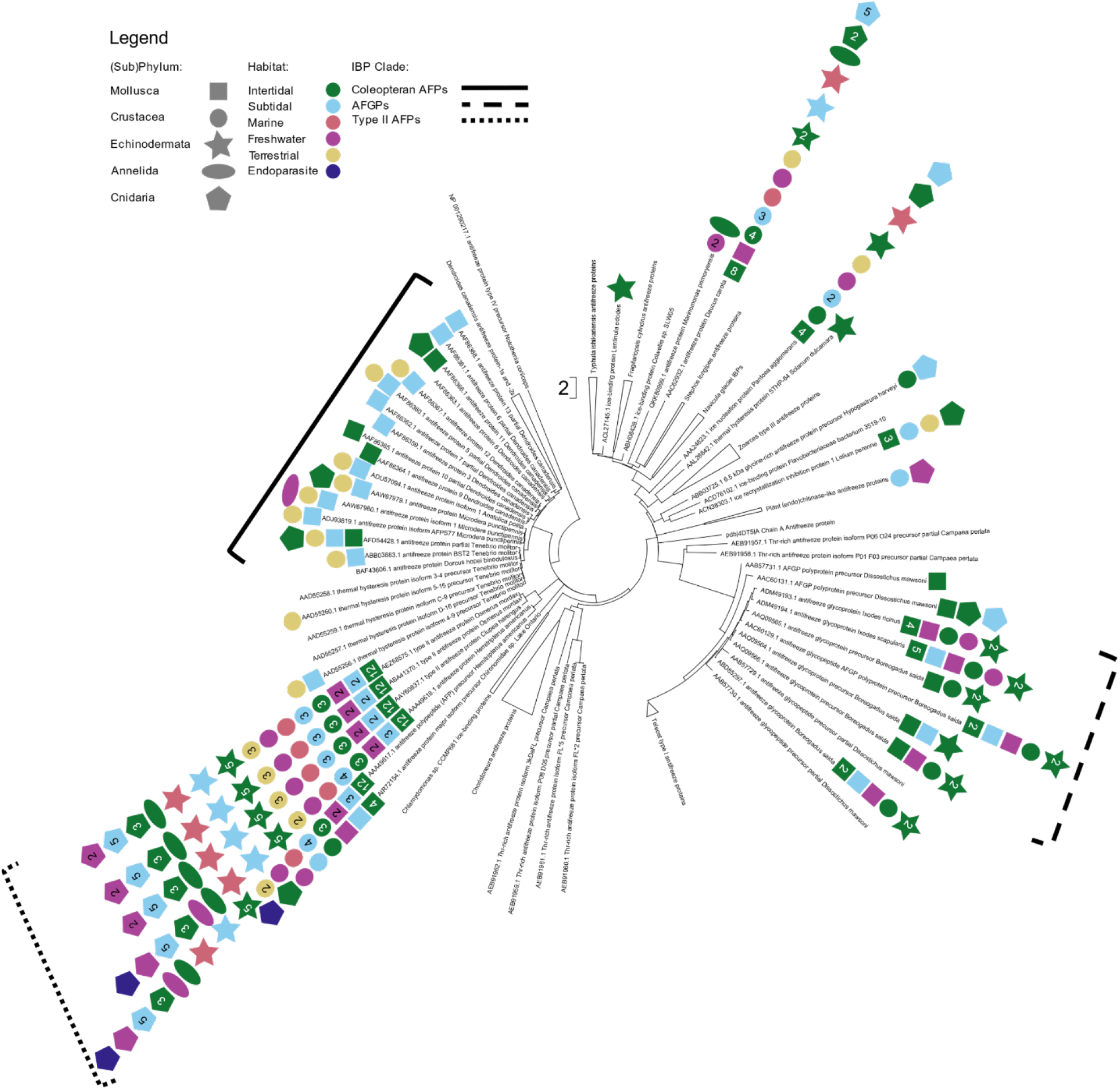
Phylogeny of query IBPs with invertebrates bearing sequences with homology to queries organized by (sub)phyla and habitat mapped against it. Brackets highlight IBP clades of interest for patterns of invertebrates with homology to IBPs. Numbers in symbols represent the number of invertebrate species for the respective phylum and habitat type with homology to the IBP the symbol is aligned with in the tree, symbols with no number have only 1 species in that category.

**Fig. 2.**
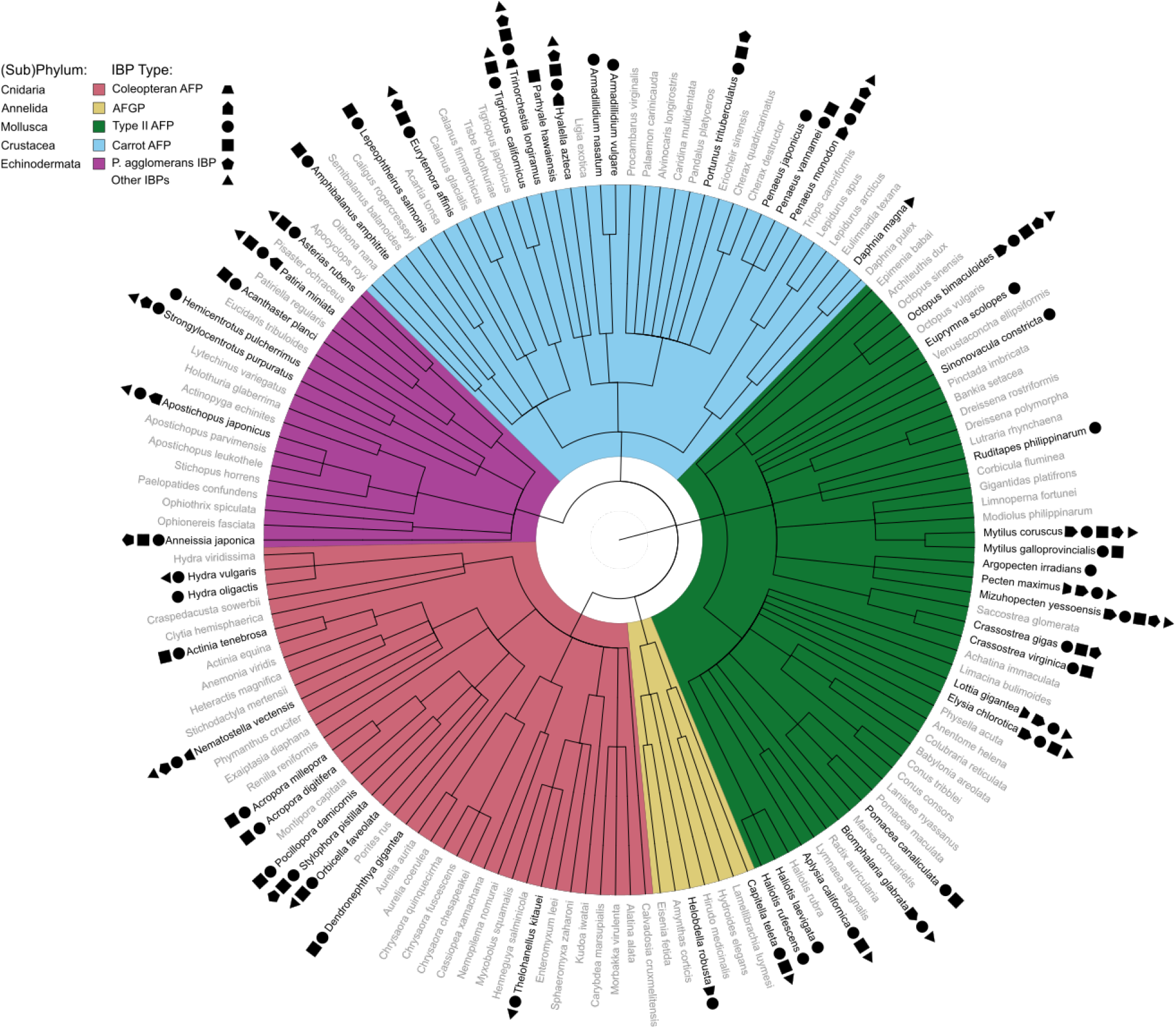
Phylogeny of invertebrate species used to search for putative IBPs and the respective query IBPs bearing high homology mapped against it, colour-coded by (sub)phylum. Black species names indicate presence of at least one putative IBP was found, the opposite is true for grey species names. Symbols represent different clades of IBPs, n _coleopteran AFP_ = 30, n _AFGP_ = 6, n _type II AFP_ = 5, n _carrot AFP_ and n _*P. agglomerans* IBP_ = 1, n _other IBPs_ = 12 sequences. Branch lengths are not informative and show overall relatedness between species determined by NCBI taxonomy.

We were interested in whether intertidal species were more likely to contain IBPs, so we investigated the relationship between the probability of IBP homology and habitat type while controlling for (sub)phylum. The probability that a species contained a protein coding sequence that had high homology to a known IBP from the query list was not impacted by the (sub)phylum of the search organism (X^2^_2,108_ = 0.93, p = 0.628; Fig. 2) nor was there an interaction between (sub)phylum and habitat (X^2^_6,102_ = 8.88, p = 0.180). However, we found that habitat type was a strong predictor of IBP homology (X^2^_3,110_ = 22.46, p < 0.001). More specifically, the proportion of intertidal invertebrates with sequence homology with known IBPs (55.3%) was significantly greater than that for freshwater (19.4%, z = 2.93, p = 0.016) and other marine species (5.0%, z = −2.93, p = 0.016), but not subtidal species (48.0%, z = −0.56, p = 0.939; Fig. 3). Subtidal invertebrates also had a higher likelihood of containing IBP homology than other marine species (z = 2.60, p = 0.042) but not freshwater species (z = 2.22, p = 0.108; Fig. 3). This probability did not significantly differ between freshwater and other marine invertebrates (z = −1.35, p = 0.512; Fig. 3).

**Fig. 3.**
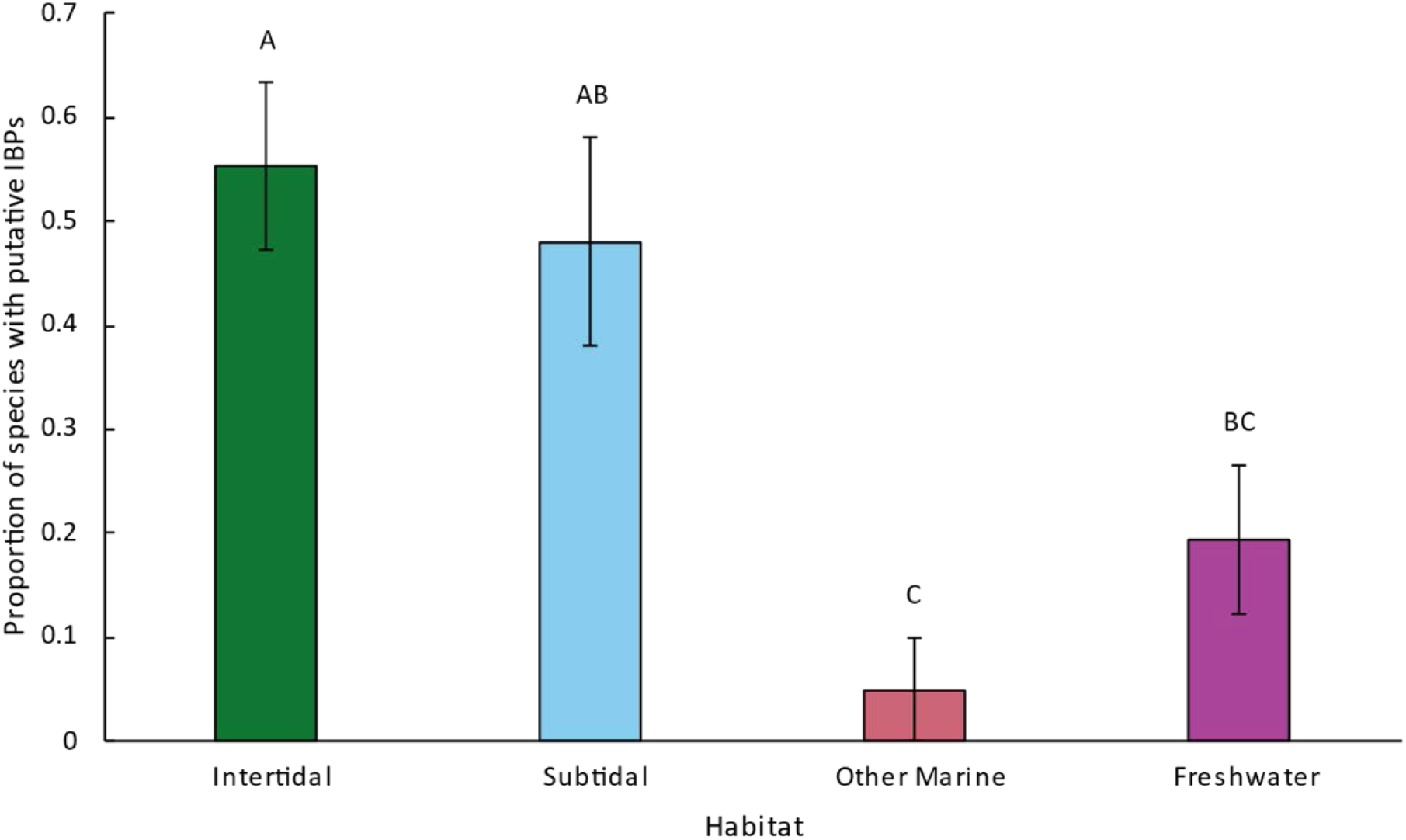
Comparison of proportion of invertebrate species bearing homology with IBPs across by their habitats. IBP homology found through a protein BLAST search with an expect threshold of 1 × 10^−5^. Letters denote statistical difference between habitats, habitats that do not bear the same letter are significantly different from each other (p < 0.05). n _intertidal_ = 38, n _subtidal_ = 25, n _other_, n _marine_ = 20, n _freshwater_ = 31. Error indicates the standard error of the proportion.

### Assessing Putative IBPs

The phylogeny of query IBPs grouped structurally similar IBPs into clades, with little evidence of phylogenetic signal; for example, tick AFGPs grouped with fish AFGPs rather than with IBPs from other arthropods, while type II AFPs from fish grouped in the middle of a largely arthropod IBP clade rather than with other fish IBPs (Fig. 1). Of the identified IBP clades, only three had significant homologies with the invertebrates in our organism search list: Coleoptera AFPs, AFGPs, and type II AFPs (Fig. 1). An AFP from a carrot (*Daucus carota*) and an ice-nucleating IBP from a bacterium (*Pantoea agglomerans*) also had strong but isolated homology with sequences from multiple search organisms (Fig. 1).

We wanted to investigate the above three clades further to ensure these trends were due to an IBP-specific search. To isolate shared sequence regions between IBPs we obtained eight ancestral sequences from the Coleoptera AFP, AFGP, and type II AFP clades: four from the Coleoptera AFP clade, two from the AFGP clade, and two from the type II AFP clade (Table 3). All ancestral sequences returned hits for IBPs in the non-redundant protein sequence database in NCBI (1988) and where applicable, did not have matches with sequences for progenitor proteins (Table 3), suggesting we successfully isolated these shared IBP-specific sequence regions. When searching for these ancestral sequences in the organism search list, two from the Coleoptera AFP clade (both from *Dendroides canadensis*) did not return hits, suggesting the initial search from these sequences were not specific for IBPs. For the other two Coleopteran AFP query ancestral sequences, one returned matches in the same two species as the original queries, but the other ancestral sequence had matches in all but two of the original five species. In the case of the ancestral sequences of the AFGP queries, three of the original seven species were missing for the fish AFGPs and two of the 11 were missing for those of the ticks. In the case of the ancestral sequences for the type II AFPs, matches were found in all the same species as the original queries. With these exceptions, this analysis suggested that our search method was specific in finding putative IBPs in our search organisms.

**Table 3.**
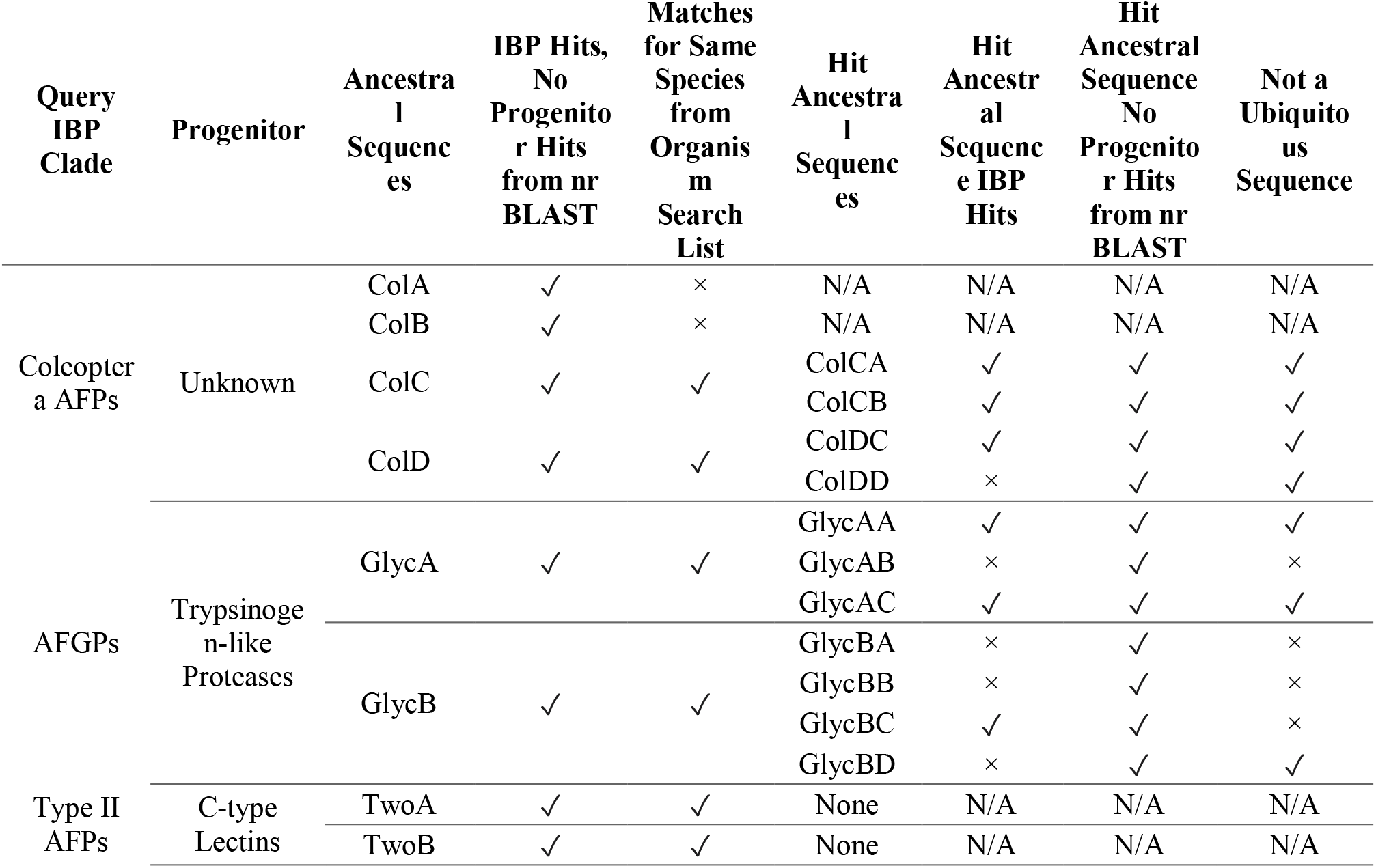
Assessment of putative IBPs through a series of ancestral sequences and BLAST searches. Query IBP clade refers to clades of IBPs identified in Fig 1. Progenitor sequence is identified from Bildanova et al. (2012). Ancestral sequences were calculated from query sequences in the respective clades. Hit ancestral sequences were calculated from the BLAST matches for the ancestral sequences in our search organism list. “IBP hits” refers to BLAST matches for IBPs, “no progenitor hits” indicates that there were no BLAST matches with IBP progenitors, and “not ubiquitous sequence” means that there was not a broad taxonomic range of BLAST matches. All BLAST searches were conducted against the entire non-redundant protein database of NCBI.

We next aimed to determine if the shared sequence regions of the matches for our above ancestral sequences in our putative IBPs were IBP-specific rather than for IBP progenitors or for other non-IBPs common across a wide range of organisms. For the matches of the Coleopteran AFP and the AFGP ancestral sequences, four and seven sequences were obtained, respectively (Table 3). For all but one of the sequences obtained through the matches for Coleopteran AFP ancestral sequences, matches for IBPs were found and, in all cases, the narrow taxonomic distribution of these matches suggested that the sequence type is not ubiquitous and may be a derived IBP (Table 3). Sequences obtained from the matches for AFGP ancestral sequences were less consistent, with only three sequences returning matches for IBPs and three sequences returning matches across a broad taxonomic range, suggesting a ubiquitous sequence and an overall less specific search for IBPs (Table 3). Despite this, no hits for the progenitor trypsinogen-like proteases were obtained. The ancestral sequences were not obtained in the case of the matches for the type II AFP ancestral sequences; for reasons unknown the phylogeny produced from these matches had multiple polytomies and could not be used to obtain ancestral sequences.

We next aimed to further assess ice-binding capabilities of the putative IBPs that we identified. Because only the top hits for each search organism were used for the type II AFPs and the carrot AFP, a total of 250 putative IBPs were examined using each of the three IBP calculators (He et al., 2015; Xiao et al., 2016; Pratiwi et al., 2017) and 86 of these sequences were also examined with a disulfide bridge calculator (Ferrè and Clote, 2005a,b, 2006). DiANNA 1.1 (Ferrè and Clote, 2005a,b, 2006) indicated that 69.8% of protein sequences with high homology to type II AFPs had five or more disulfide bridges and identified all query type II AFPs sequences as having five or more disulfide bridges. TargetFreeze (He et al., 2015) calculated 60.0% of the sequences to be IBPs, compared to 49.6% from iAFP-Ense (Xiao et al., 2016), and 65.2% from CryoProtect (Fig. 4; Pratiwi et al., 2017). This meant that while on average each calculator rejected 41.7% of the sequences, only 17.6% of the putative IBP sequences were not identified as IBPs by any calculator. When vetting the 42 query sequences with high homology to search organism sequences, TargetFreeze (He et al., 2015) accepted only 73.8% of the query sequences (Fig. 4), with all query AFGPs rejected. By contrast, iAFP-Ense (Xiao et al., 2016) accepted 90.5% of the query sequences and CryoProtect (Pratiwi et al., 2017) accepted 95.2% (Fig. 4). This means each calculator rejected an average of 13.5% query sequences, but only one query IBP (2.4%), a thermal hysteresis protein from bittersweet nightshade (*Solanum dulcamara*; NCBI accession: AAL26842.1) was rejected by all three IBP calculators. In many cases, including the AFGPs for TargetFreeze (He et al., 2015), a query sequence getting rejected by the calculators did not equate to all respective hits getting rejected as well (Table S4).

**Fig. 4.**
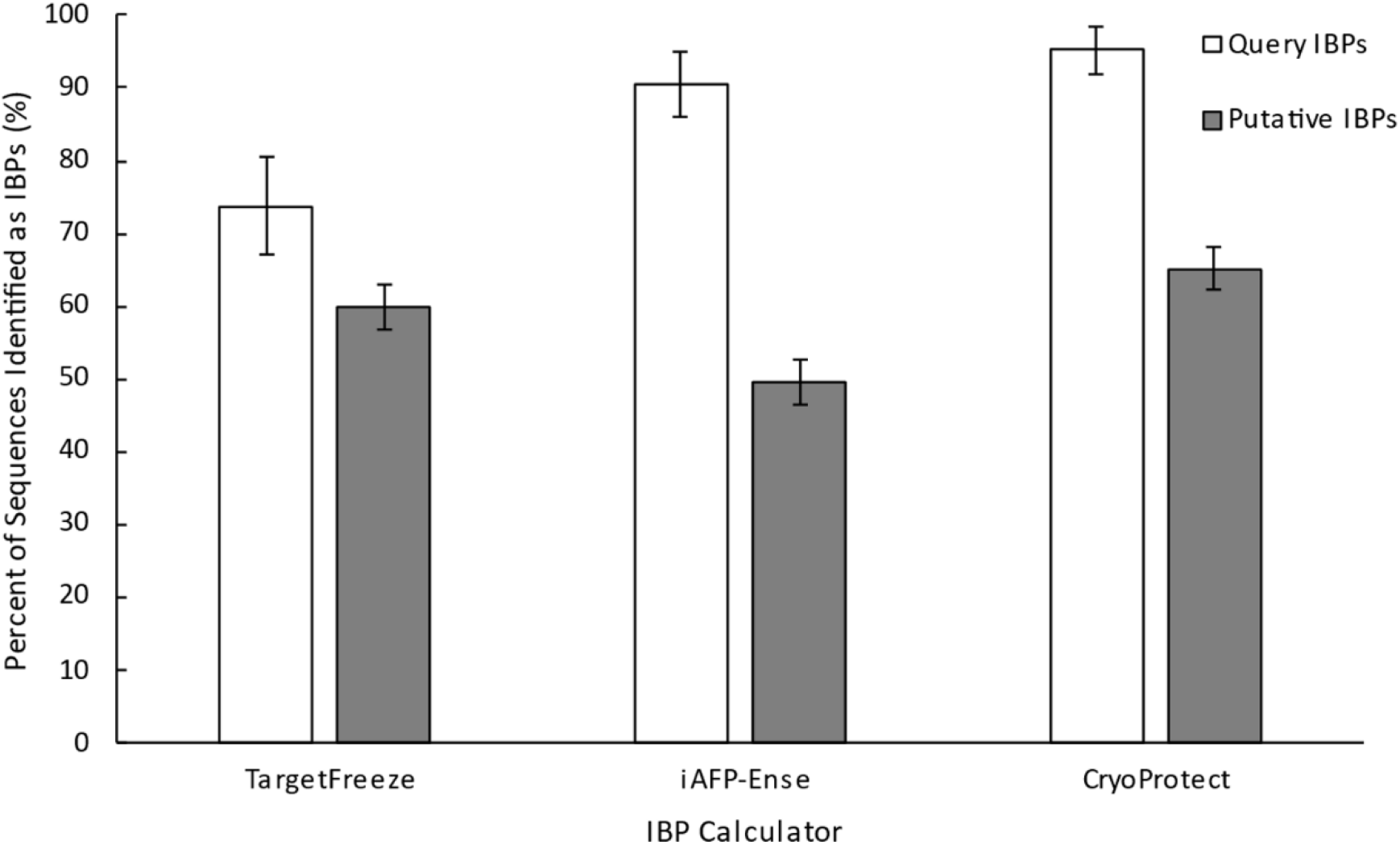
The percentage of query IBP sequences (n = 42) and putative IBP sequences (n = 250) identified as IBPs by three IBP calculators: TargetFreeze (He et al., 2015), iAFP-Ense (Xiao et al., 2016), and CryoProtect (Pratiwi et al., 2017). Error bars represent the standard error of the proportion.

### Investigating Evolutionary Origin of a Putative IBP

To evaluate the evolutionary origin of a putative IBP from the hard-shelled mussel (*Mytilus coruscus*), we evaluated the level of synteny in a group of search organisms with well-developed genomic resources and potential orthologs of the putative IBP (NCBI accession: CAC5422424.1; Fig. 4). We selected the four nearest genes to the coding region of the *M. coruscus* putative IBP: two upstream and two downstream, referred to as gene or protein 1, 2 and 4, 5 respectively henceforth (NCBI accessions: CAC5422422.1, CAC5422423.1, CAC5422425.1, CAC5422426.1). Interestingly, gene 2 had high homology to the *M. coruscus* putative IBP, with 100% query coverage, 70% identity, and an e-value of 2 × 10^−91^, suggesting that these two genes may be the result of a gene duplication event or, less likely, a technical artifact of the genome assembly process. No evidence for synteny across these five genes was found in any of the species, regardless of relatedness to *M. coruscus* (Fig. 5A, B). In nearly all species, the presumed ortholog for the putative IBP was isolated, with no orthology for the surrounding genes on the same contig, scaffold or even chromosome save gene 2, which would have potential orthologs of varying quality directly overlapping the orthologs of our putative IBP gene. The one exception was the great scallop (*Pecten maximus*), where gene 5 was found on the same chromosome as the ortholog for the putative IBP but located millions of base pairs away from it, suggesting that this relationship does not reflect true synteny at our level of interest.

**Fig. 5.**
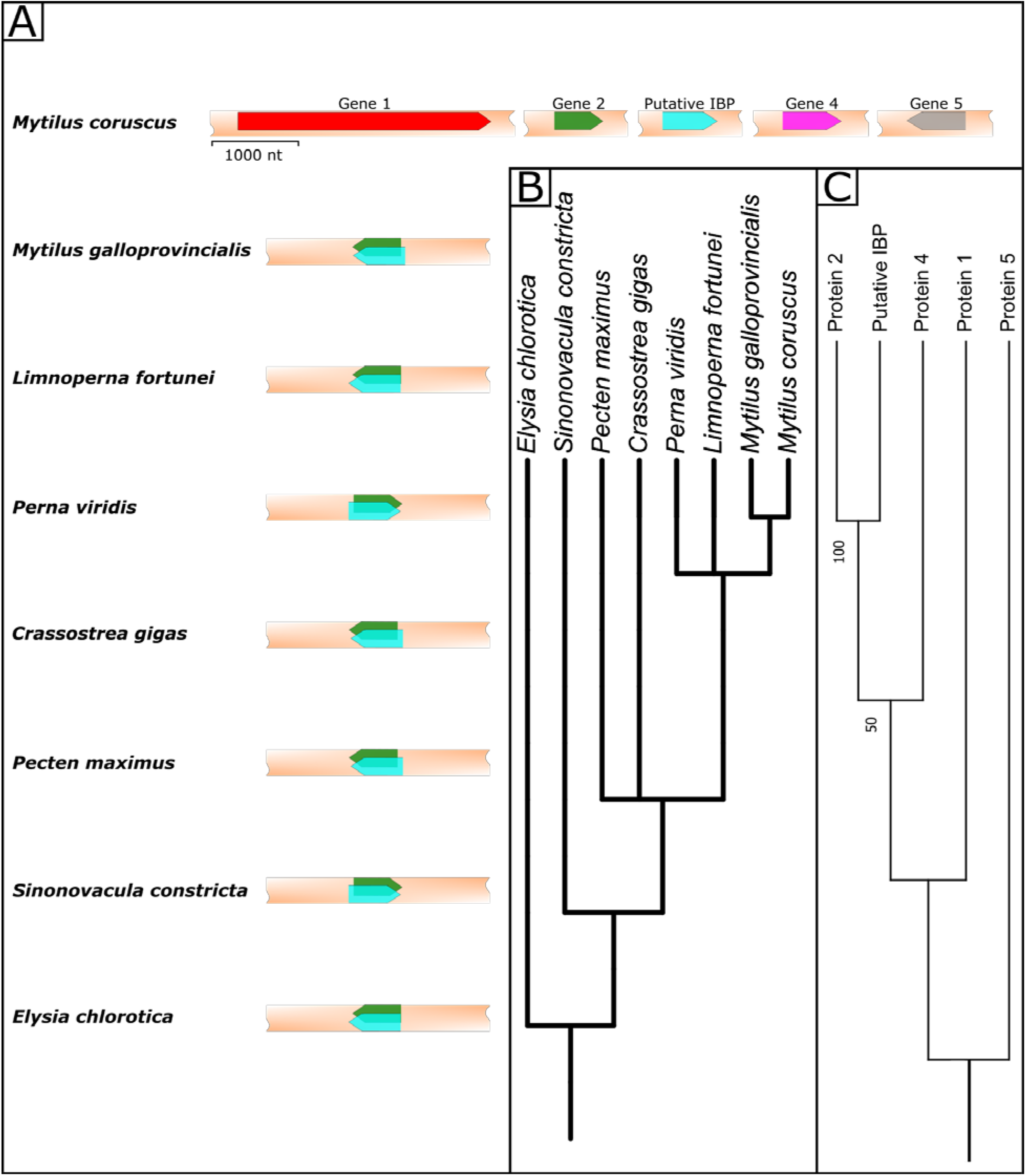
Evolution of a putative IBP in *M. coruscus*. A) Arrangement of the coding region for the putative IBP of *Mytilus coruscus* and its 4 surrounding genes relative to their predicted orthologs in other mollusc species. B) Phylogeny of the mollusc species observed, demonstrating degree of relatedness to each other. **C)** Phylogeny of the 5 *Mytilus coruscus* proteins from the coding regions shown in panel “A”, demonstrating degree of similarity between sequences.

In the Mediterranean mussel (*Mytilus galloprovincialis*), all top candidates for orthologs of the putative IBP were in the exact same locations as the potential orthologs for gene 2, but E-values were lower and percent identities were greater for gene 2 compared to those for our putative IBP suggesting the presence of a gene 2 ortholog, rather than one for the putative IBP. This overlap in ortholog sites for gene 2 and the putative IBP is also seen in the golden mussel (*Limnoperna fortunei*) and *Elysia chlorotica* a sea slug, but in the latter the potential ortholog for the putative IBP outscores that of gene 2. In all cases, BLAST output alignments had overlap of varying scores between potential ortholog sites for the putative IBP and gene 2, matching the aforementioned high similarity between sequences (Fig. 5A). This similarity in sequences is reflected in their grouping and branch lengths, representing number of substitutions, in the phylogeny produced from our protein of interest and the 4 surrounding proteins (Fig. 5C). In addition to this, the scores for the orthologs for both gene 2 and the putative IBP drop in power after the most closely related species *M. galloprovincialis* by well over 10 orders of magnitude.

## DISCUSSION

Here we provide evidence that IBPs evolve relatively easily in response to the risk of freezing. We found dozens of invertebrate species with sequences with high homology to known IBPs, supporting our hypothesis that IBPs occur across a much broader array of organisms than currently known. We predicted that the freezing risk in intertidal habitats would select for IBP presence and found much higher probabilities of putative IBP presence in intertidal species than species found in habitats that have lower risk of freezing. These findings strongly suggest that there are uncharacterized IBPs in intertidal invertebrates that have high homology to teleost type II AFPs and AFGPs supporting our hypothesis that IBPs are selected for by intertidal habitats.

This is not the first study to use molecular sequence data to predict the presence of IBPs (Krell et al., 2008), but this is the first to our knowledge taking this broad approach. Typically, IBPs are predicted from sequence data of specific species during genome annotation projects or transcriptomics research. For example, the genome project for the scallop *Mizuhopecten yessoensis* resulted in 15 predicted protein products annotated as ice-nucleating proteins alone (Wang et al., 2017). In these other studies, IBP presence is predicted by homology to identify an uncharacterized coding region or transcript, thus filling gaps in our knowledge for individual species. By contrast, in this study we aimed to fill gaps in our knowledge of IBP evolution by taking a broad approach to identify putative IBPs and identify potential correlations with IBP abundance and specific habitat types. A major limitation of this study that is shared with genome and transcriptome studies is the inability to confirm the expression or ice-binding activity of these proteins. An advantage of our study and its broad search list is the potential to avoid false negatives that might arise due to limited queries in traditional automated gene annotation processes. This could result in the under-prediction of a given protein type, a phenomenon that is more common in the case of functionally rare and unique genes such as IBPs (Prosdocimi et al., 2012; Wilbrandt et al., 2019).

Despite the above limitations, validation of our search process suggests some degree of specificity for identifying IBPs rather than matches due to sequences unrelated to IBP activity, with some exceptions noted below. Our initial verification using a BLAST search of the queries and their ancestral sequences beyond our search organisms and against all NCBI’s non-redundant database yielded results for IBPs, not their progenitor proteins. These query ancestral sequences also yielded hits in nearly all the same species as the initial IBPs save for a select group of Coleopteran AFPs. We also attempted to verify our search method using the ancestral sequences of the matches, which suggested that our search method may not be specific in the case of some of the AFGPs as most of the sequences did not return matches for IBPs and returned matches across a wide taxonomic range. However, in all this suggests that our search process was specific for finding derived IBPs and not general similarities in sequences that are coincidentally more common in intertidal invertebrates. Also, given the nature of how expect thresholds are calculated and implemented in determining search results in BLAST, it is possible more potential IBPs exist but were rejected (González-Pech et al., 2019). In fact, other than being unable to isolate and characterize these proteins, all limitations of this study will likely lead to underestimation of true IBP frequency. For example, the advent of shotgun sequencing means genomic resource availability is constantly increasing, expanding the search potential for IBPs (Giani et al., 2020). Similarly, there are a broad array of IBPs that have been characterized in the literature, but not yet sequenced (Madison et al., 1991; Duman et al., 2004). As reflected by the IBP sequence phylogeny (Fig. 1), IBP types do not have a common sequential or structural component that unites them all and it is probable that new structures of IBPs have yet to be discovered that would expand our search potential even further (Davies, 2014; Bar Dolev et al., 2016).

Due to this lack of shared sequence or structure across IBPs, there has been extensive work in the field of machine learning to identify IBPs from their amino acid sequence (e.g. Kandaswamy et al., 2011; He et al., 2015; Yang et al., 2015; Xiao et al., 2016; Pratiwi et al., 2017; Eslami et al., 2018; Khan et al., 2018; Nath and Subbiah, 2018; Usman and Lee, 2019; Sun et al., 2020; Usman et al., 2020; Wang et al., 2021). We used three IBP calculators in this study to further investigate our putative IBPs. Although most of the sequences identified by our BLAST searches as putative IBPs passed as IBPs according to at least one calculator, on average each calculator rejected 41.7% of the sequences. These calculators were created because IBPs are more similar to their progenitor non-IBPs than each other, so the large fraction of proteins rejected by each calculator is unsurprising (Kandaswamy et al., 2011; Eslami et al., 2018; Nath and Subbiah, 2018; Sun et al., 2020; Usman et al., 2020). What is surprising however is the variability among calculator outputs. Despite each calculator rejecting an average of 41.7% of the putative IBPs, only 17.6% of the sequences were rejected by all calculators. Similarly, an average of 13.5% of query sequences were also rejected by each calculator, but only one was rejected by all three, suggesting that these calculators also struggle to predict IBPs individually, but we can have more confidence where the three overlap in their rejections. It also demonstrates the importance of training and testing datasets in creating machine learning based IBP predictors, especially in the case of TargetFreeze (He et al., 2015) which rejected all query AFGPs, indicating a clear gap in the algorithm. By contrast, all calculators recognized the query carrot IBP but rejected nearly all its homologous sequences from the species search list. This may be an instance where the IBP calculators were powerful in their ability to differentiate IBPs from non-IBPs. It should be noted however, that this carrot query IBP is quite unique even compared to other plant IBPs (Wang et al., 2020) and it may be that the calculators were trained with the sequence, meaning it could correctly identify it, but since it is unique in the database of IBPs the calculators were not sufficiently trained in identifying other carrot-like IBPs, despite these IBP calculators being designed taking into account the impact of unbalanced data (Murphey and Guo, 2004; Usman et al., 2020). Expanded empirical studies of ice-binding activities will be necessary to resolve these uncertainties.

We also used DiANNA 1.1, the disulfide bridge calculator, to verify a subset of our hits for the type II AFPs (Ferrè and Clote, 2005a,b, 2006). DiANNA 1.1 is mainly intended for identifying the potential locations of disulfide bridges, rather than the amount, meaning it often overestimates the number of disulfide bridges (Ferrè and Clote, 2005a,b, 2006). Despite this, the calculator providing outputs for fewer than 5 disulfide bridges still acted as a sufficient criterion for rejection as in Graham et al. (2008). There are IBP calculators that exist that are precise in the identification of IBPs from C-type lectins, but these have prohibitive computational requirements (Kozuch et al., 2018). These highlighted shortcomings in the *in silico* research of IBPs emphasizes the value of experimental verification of ice-binding activity.

The limited laboratory work that exists on IBPs in intertidal invertebrates also suggest they may be proteins with similarities to known AFGPs and type II AFPs in these species. The earliest instance of IBP identification in an intertidal invertebrate was an AFGP in the blue mussel (*Mytilus edulis*; Theede et al., 1976). Here we found many intertidal species bearing strong homologous sequences to both teleost and tick AFGPs; however, *M. edulis* does not have a sequenced genome and was therefore absent from our search list, and so this could not be replicated in this study. *Mytilus* species that were included in our organism list did not have homologous sequences to AFGPs (save *M. coruscus* which had an unnamed protein product with homology to one of the tick AFGPs) but other molluscs had sequences with strong homology to the query AFGPs (Fig. 1). It is unknown how much homology, if any, the *M. edulis* AFGP has with the tick and teleost AFGPs and it should be noted that while observation of ice-binding activity has since been repeated in *M. edulis* (Dubé, 2012), no characterization of the actual protein has been performed since the original study in 1976 (Theede et al., 1976; Duman, 2015). With respect to type II AFPs, actual isolation of a protein with high homology to this structure has not been completed in an intertidal invertebrate. However, a transcriptomic study comparing transcription before and after freezing in an intertidal barnacle (*Semibalanus balanoides*) found many transcripts automatically annotated as macrophage mannose receptors upregulated (Marshall et al., 2018 preprint). Mannose receptors are not associated with freeze tolerance, but other C-type lectins, specifically type II AFPs, are clearer contributors to freeze tolerance (Gronwald et al., 1998). It was hypothesized that these upregulated transcripts were misannotated and if this were the case it would corroborate the results of this study that found a strong signal for type II AFPs in intertidal invertebrates (Marshall et al., 2018 preprint). It should be noted however, *S. balanoides* was in our search organism list, but we did not find any strong homology to query IBPs.

There were also many genomic matches with a carrot (*Daucus carota*) IBP and bacterium (*Pantoea agglomerans*) IBP. The structure of the carrot IBP is quite unique in sharing an ice-binding site structure with grass AFPs but otherwise having an irregular arrangement of surface residues not seen in other IBPs, and the calculators suggest many of these hits may not be IBPs (Wang et al., 2020). The bacterial IBP was specifically one with ice nucleating activity (Warren and Corotto, 1989). Ice nucleating proteins have been found in intertidal invertebrates before, specifically an air breathing gastropod (*Melampus bidentatus*) but it is unclear how homologous they would be to those from bacteria (Madison et al., 1991). The *M. bidentatus* ice-nucleating proteins were found to have similar amino acid proportions as bacterial ice-nucleating proteins, but the size of the protein was only a quarter of those found in bacteria and the sequence was not obtained (Madison et al., 1991). Given the size difference, it is unlikely the results of our study reflect the presence of a *M. bidentatus*-like IBP distributed across intertidal invertebrates. In the intertidal bivalve (*Geukensia demissa*), freeze tolerance is accomplished through ice-nucleating bacteria, rather than proteins produced by the animal itself (Loomis and Zinser, 2001). Whether a similar relationship is occurring in multiple other intertidal invertebrates is unknown, but considering how ubiquitous *P. agglomerans* is, and how frequently it acts as a symbiont, bacterial contamination of samples could partially account for the high signal for this IBP (Dutkiewicz et al., 2016).

While the Coleoptera AFP clade did not have a strong intertidal signal like the previously mentioned IBP groups, multiple query IBPs in this clade had homology for a protein labelled as an “insect AFP” in a terrestrial crustacean (*Trinorchestia longiramus*). As with the other hits in this study, this is a predicted AFP from a genome annotation project, not a laboratory-characterized AFP (Patra et al., 2020). Given how closely insects and crustaceans are related compared to the other search phyla and the taxonomy of the query IBPs with strong signals, potential evolutionary homology could explain the acquisition of this crustacean’s putative IBP. This may also be true for the strong homologies found for the tick AFGPs in some crustaceans in the search organism list, but none were automatically annotated as IBPs in these cases. Despite how closely related insects and ticks are to crustaceans, nothing from the query list is more closely related to crustaceans than other crustaceans. The query IBPs from a copepod (*Stephos longipes*) did not contain homologies to any sequences in the search organisms, when we initially expected this protein may return multiple hits in the crustaceans in our search list. These proteins are hypothesized to be acquired through lateral gene transfer from diatoms or snow molds, both groups of IBPs that also did not yield hits in this study, which explains why no hits were found in these query IBPs despite being part of the search phyla (Kiko, 2010). Lateral gene transfer has been suggested for multiple other IBPs as well, with evidence for IBP acquisition through lateral gene transfer being seen in algae, diatoms, and fungi (Sorhannus, 2011; Raymond, 2014; Arai et al., 2019; Raymond and Remias, 2019).

Type II AFPs in some fish have also been hypothesized to have been acquired through lateral gene transfer (Graham et al., 2012). Unlike previously-mentioned examples of IBP lateral gene transfer, the type II AFPs remain confined within the same phylum, making it unlikely that lateral gene transfer explains the strong signal seen for type II AFPs seen in this study. This is corroborated by the fact that none of the hits found in this study were more than 60% identical to the respective query IBP sequences, despite the high homology suggested by hits surpassing the expect threshold of 1 × 10^−5^. This suggests that all IBPs in the search organisms of this study were acquired convergently, including those for crustaceans highlighted in the previous paragraph. Type II AFPs evolved by gene duplication and neofunctionalization from C-type lectins in fish, and it is possible a similar process occurred in the C-type lectins of these intertidal taxa (Gronwald et al., 1998). The same can also be hypothesized for AFGPs, as they have evolved convergently within fish, both from trypsinogen-like proteases and non-sense DNA. A similar evolutionary pathway could explain the putative presence of AFGPs in our search organisms (Chen et al., 1997; Zhuang et al., 2019). Further research into the potential evolution of these IBPs could provide further understanding of the colonization of the intertidal zone in temperate and polar regions.

We investigated the potential evolutionary origin of these putative IBPs by selecting a strong IBP match in a search organism, *Mytilus coruscus*, with many closely related species with genomic resources to be treated as a proof of principle. A basic search for synteny proved fruitless in this study, which can be used as support for horizontal gene transfer (Sevillya et al., 2020). However, for reasons mentioned above, we believe that this IBP evolved convergently and horizontal gene transfer could not be responsible for the acquisition of this protein. Despite this lack of synteny being found, our search methods revealed a high similarity between the potential IBP of *Mytilus coruscus* and an unnamed protein product with a neighbouring coding region. Based on potential orthologs, we also found that neither of these two proteins are very conserved beyond *Mytilus galloprovincialis*, which appeared to only have the neighbouring protein and not an ortholog for the potential IBP. Based on this, we suggest a duplication and neofunctionalization event occurred with this neighbour protein resulting in an IBP in *Mytilus coruscus*. This duplication and neofunctionalization would mirror the hypothesized evolutionary history for other known IBPs including fish type II AFPs (Liu et al., 2007) and type III AFPs (Deng et al., 2010), suggesting that evolutionary history repeated itself in intertidal invertebrates. However, gene 2 and its ortholog in *M. galloprovincialis* may be putative IBPs themselves that evolved *de novo* as seen in certain fish AFGPs (Zhuang et al., 2019), based on it not being conserved beyond the genus *Mytilus*. This would mean a duplication event occurred creating paralogs that both function as IBPs in *M. coruscus*, this may reflect the origin of the many AFP paralogs seen in some fish and insects (Swanson and Aquadro, 2002). The evolutionary origin of IBPs is still a developing field, with the evolution of most non-fish IBPs poorly studied, but there are also examples of IBPs obtained convergently in the case of AFGPs and type I AFPs from fish (Chen et al., 1997; Graham et al., 2013). Convergent evolution of similarly structured IBPs across phyla has not been explicitly described in the literature before, as most known instances of similar IBPs between phyla have been explained by horizontal gene transfer (Kiko, 2010; Sorhannus, 2011; Raymond, 2014; Arai et al., 2019; Raymond and Remias, 2019). This would make our proposed convergent evolution of this potential type II AFP-like IBP in *Mytilus coruscus* unique. However, given the number of putative IBPs and their taxonomic distribution (Fig. 2), it is likely there are many other examples yet to be described. This combined with the multitude of confirmed IBPs suggests that convergent evolution of IBPs is common.

In this study, we demonstrate evidence for an enrichment of uncharacterized putative IBPs in intertidal invertebrates as compared to invertebrates living in other aquatic habitats, supporting the hypothesis that life in the intertidal zone might select for IBPs. In intertidal invertebrates, a strong signal for putative IBPs was found for type II AFPs from fishes and AFGPs from both fishes and ticks, reflecting existing data of AFGPs in intertidal mussels and evidence of type II AFPs in intertidal barnacles. We suggest that these IBPs evolved convergently, potentially through a duplication and neofunctionalization event as we suggest for *M. coruscus*. Future studies will be necessary to confirm the expression and true ice binding activity of these proteins, but we were able to demonstrate both through reverse BLAST searches and IBP calculators that our methods were specific in their approach, suggesting we detected IBPs and not simply proteins with high similarity to IBPs. Even with rejections considered from our alternate BLAST searches and IBP calculators, the trends seen in habitat specific IBP presence holds true. We therefore conclude that IBPs readily evolve in response to intertidal environments, and this could help explain how intertidal organisms colonized temperate and polar regions.

## Acknowledgements

We would like to thank Patricia Schulte and Jeffrey Richards for their input which helped develop our methods and interpret our results. We used FigTree in creating one of our figures, created and made available to the public by Andrew Rambaut.

## Data availability

All data is available on the Open Science Framework (link will be replaced with DOI on acceptance): https://osf.io/b7a92/?view_only=b8648c8a3d114438a6fdbe70a20c5cca

## REFERENCES

Aarset, A. V. (1982). Freezing tolerance in intertidal invertebrates (a review). Comp. Biochem. Physiol. 73A, 571–580.

Altschul, S. F., Gish, W., Miller, W., Myers, E. W. and Lipman, D. J. (1990). Basic local alignment search tool. J. Mol. Biol. 215, 403–410.

Arai, T., Fukami, D., Hoshino, T., Kondo, H. and Tsuda, S. (2019). Ice-binding proteins from the fungus Antarctomyces psychrotrophicus possibly originate from two different bacteria through horizontal gene transfer. FEBS J. 286, 946–962.

Arning, N. (2018). Ancestral sequence reconstruction. DTC Coding Dojo. Doctoral Training Centre University of Oxford. Oxford, UK. URL: https://dtc-coding-dojo.github.io/main//blog/Ancestral_sequence_reconstruction/.

Balcerzak, A. K., Capicciotti, C. J., Briard, J. G. and Ben, R. N. (2014). Designing ice recrystallization inhibitors: from antifreeze (glycol)proteins to small molecules. RSC Adv. 4, 42682–42696.

Bar Dolev, M., Braslavsky, I. and Davies, P. L. (2016). Ice-binding proteins and their function. Annu. Rev. Biochem. 85, 515–542.

Bildanova, L. L., Salina, E. A. and Shumny, V. K. (2012). Main properties and evolutionary features of antifreeze proteins. Russ. J. Genet. Appl. Res. 3, 66–82.

Chen, L., DeVries, A. L. and Cheng, C.-H. C. (1997). Evolution of antifreeze glycoprotein gene from a trypsinogen gene in Antarctic notothenioid fish. Proc. Natl. Acad. Sci. USA, 94, 3811–3816.

Davies, P. L. (2014). Ice-binding proteins: a remarkable diversity of structures for stopping and starting ice growth. Trends Biochem. Sci. 39, 548–555.

Deng, C., Cheng, C.-H. C., Ye, H., He, X. and Chen, L. (2010). Evolution of an antifreeze protein by neofunctionalization under escape from adaptive conflict. PNAS 107, 21593–21598.

Dubé, A. (2012). Investigation of antifreeze protein activity in blue mussels and amyloid-like transition in a predominant winter flounder serum antifreeze protein. MSc Thesis, Dalhousie University, Halifax, NS.

Duman, J. G. (2015). Animal ice-binding (antifreeze) proteins and glycolipids: an overview with emphasis on physiological function. J. Exp. Biol. 218, 1846–1855.

Duman, J. G., Bennett, V., Sformo, T., Hochstrasser, R. and Barnes, B. M. (2004). Antifreeze proteins in Alaskan insects and spiders. J. Insect Physiol. 50, 259–266.

Dutkiewicz, J., Mackiewicz, B., Lemieszek, M. K., Golec, M. and Milanowski, J. (2016). Pantoea aglomerans: a mysterious bacterium of evil and good. Part IV. Beneficial effects. Ann. Agric. Environ. Med. 23, 206–222.

Edgar, R. C. (2004). MUSCLE: Multiple sequence alignment with high accuracy and high throughput. Nucleic Acids Res. 32, 1792–1797.

Eslami, M., Zade, R. S. H., Takalloo, Z., Mahdevar, G., Emamjomeh, A., Sajedi, R. H. and Zahiri, J. (2018). afpCOOL: A tool for antifreeze protein prediction. Heliyon 4, e00705.

Ferrè, F. and Clote, P. (2005a). DiANNA: a web server for disulfide connectivity prediction. Nucleic Acids Res. 33, W230–W232.

Ferrè, F. and Clote, P. (2005b). Disulfide connectivity prediction using secondary structure information and diresidue frequencies. Bioinform. 21, 2336–2346.

Ferrè, F. and Clote, P. (2006). DiANNA 1.1: an extension of the DiANNA web server for ternary cysteine classification. Nucleic Acids Res. 34, W182–W185.

Fletcher, G. L., Hew, C. L. and Davies, P. L. (2001). Antifreeze proteins of teleost fishes. Annu. Rev. Physiol. 63, 359–390.

Giani, A. M., Gallo, G. R., Gianfranceschi, L. and Formenti, G. (2020). Long walk to genomics: History and current approaches to genome sequencing and assembly. Comput. Struct. Biotechnol. J. 18, 9–19.

González-Pech, R. A., Stephens, T. G. and Chan, C. X. (2019). Commonly misunderstood parameters of NCBI BLAST and important considerations for users. Bioinformatics, 35, 2697–2698.

Graham, L. A., Lougheed, S. C., Ewart, K. V. and Davies, P. L. (2008). Lateral transfer of a lectin-like antifreeze protein gene in fishes. PLoS ONE 3, e2616.

Graham, L. A., Li, J., Davidson, W. S. and Davies, P. L. (2012). Smelt was the likely beneficiary of an antifreeze gene laterally transferred between fishes. BMC Evol. Biol. 12, 190.

Graham, L. A., Hobbs, R. S., Fletcher, G. L. and Davies, P. L. (2013). Helical antifreeze proteins have independently evolved in fishes on four occasions. PLoS ONE 8, e81285.

Gronwald, W., Loewen, M. C., Lix, B., Daugulis, A. J., Sönnichsen, F. D., Davies, P. L. and Sykes, B. D. (1998). The solution structure of type II antifreeze protein reveals a new member of the lectin family. Biochemistry, 37, 4712–4721.

He, X., Han, K., Hu, J., Yan, H., Yang, J.-Y., Shen, H.-B. and Yu, D.-J. (2015). TargetFreeze: identifying antifreeze proteins via a combination of weights using sequence evolutionary information and pseudo amino acid composition. J. Membr. Biol. 248, 1005–1014.

Henikoff, S. and Henikoff, J. G. (1992). Amino acid substitution matrices from protein blocks. Proc. Natl. Acad. Sci. USA 89, 10915–10919.

Horton, T., Kroh, A., Ahyong, S., Bailly, N., Boyko, C. B., Brandão, S. N., Gofas, S., Hooper, J. N. A., Hernandez, F., Holovachov, O. et al. (2021). World register of marine species. Flanders Marine Institute. Ostend, Belgium. URL: https://www.marinespecies.org/.

Hothorn, T., Bretz, F. and Westfall, P. (2008). Simultaneous interference in general parametric models. Biom. J. 50, 346–363.

Jia, Z., DeLuca, C. I., Chao, H. and Davies, P. L. (1996). Structural basis for the binding of a globular antifreeze protein to ice. Nature 384, 285–288.

John, U. P., Polotnianka, R. M., Sivakumaran, K. A., Chew, O., Mackin, L., Kuiper, M. J., Talbot, J. P., Nugent, G. D., Mautord, J., Schrauf, G. E. and Spangenberg, G. C. (2009). Ice recrystallization inhibition proteins (IRIPs) and freeze tolerance in the cryophilic Antarctic hair grass Deschampsia antarctica E. Desv. Plant Cell Environ. 32, 336–348.

Jones, D. T., Taylor, W. R. and Thornton, J. M. (1992). The rapid generation of mutation data matrices from protein sequences. CABIOS 8, 275–282.

Kandaswamy, K. K., Chou, K.-C., Martinetz, T., Möller, S., Suganthan, P. N., Sridharan, S. and Pugalenthi, G. (2011). AFP-Pred: A random forest approach for predicting antifreeze proteins from sequence-derived properties. J. Theor. Biol. 270, 56–62.

Kennedy, J. R., Harley, C. D. G. and Marshall, K. E. (2020). Drivers of plasticity in freeze tolerance in the intertidal mussel Mytilus trossulus. J. Exp. Biol. 223, jeb233478.

Khan, S., Naseem, I., Togneri, R. and Bennamoun, M. (2018). RAFP-Pred: robust prediction of antifreeze proteins using localized analysis of n-peptide compositions. IEEE/ACM Trans. Comput. Biol. Bioinform. 15, 244–250.

Kiko, R. (2010). Acquisition of freeze protection in a sea-ice crustacean through horizontal gene transfer? Polar Biol. 33, 543–556.

Kozuch, D. J., Stillinger, F. H. and Debenedetti, P. G. (2018). Combined molecular dynamics and neural network method for predicting protein antifreeze activity. PNAS 115, 13252–13257.

Krell, A., Beszteri, B., Dieckmann, G., Glöckner, G., Valentin, K. and Mock, T. (2008). A new class of ice-binding proteins discovered in a salt-stress-induced cDNA library of the psychrophilic diatom Fragilariopsis cylindrus (Bacillariophyceae). Eur. J. Phycol. 43, 423–433.

Kumar, S., Stecher, G., Li, M., Knyaz, C. and Tamura, K. (2018). MEGA X: Molecular evolutionary genetics analysis across computing platforms. Mol. Biol. Evol. 35, 1547–1549.

Lee, R. E. (2010). A primer on insect cold-tolerance. In Low Temperature Biology of Insects, pp. 3–34. Cambridge: Cambridge University Press.

Ling, M. L., Wex, H., Grawe, S., Jakobsson, J., Löndahl, J., Hartmann, S., Finster, K., Boesen, T. and Šantl-Temkiv, T. (2018). Effects of ice nucleation protein repeat number and oligomerization level on ice nucleation activity. J. Geophys. Res. Atmos. 123, 1802–1810.

Liu, Y., Li, Z., Lin, Q., Kosinski, J., Seetharaman, J., Bujnicki, J. M., Sivaraman, J. and Hew, C.-L. (2007). Structure and evolutionary origin of Ca^2+^-dependent herring type II antifreeze protein. PLoS ONE 2, e548.

Loomis, S. H. and Zinser, M. (2001). Isolation and identification of an ice-nucleating bacterium from the gills of the intertidal bivalve mollusc Guekensia demissa. J. Exp. Mar. Biol. Ecol. 261, 225–235.

Madison, D. L., Scrofano, M. M., Ireland, R. C. and Loomis, S. H. (1991). Purification and partial characterization of an ice nucleator protein from the intertidal gastropod, Melampus bidentatus. Cryobiology 28, 483–490.

Marshall, K. E., Dowle, E. J., Petrunina, A., Kolbasov, G. and Chan, B. K. K. (2018). Transcriptional dynamics following freezing stress reveal selection for mechanisms of freeze tolerance at the poleward range margin in the cold water intertidal barnacle Semibalanus balanoides. Biorxiv http://dx.doi.org/10.1101/449330.

Murphey, Y. L. and Guo, H. (2004). Neural learning from unbalanced data. Appl. Intell. 21, 117–128.

Nath, A. and Subbiah, K. (2018). The role of pertinently diversified and balanced training as well as testing data sets in achieving the true performance of classifiers in predicting the antifreeze proteins. Neurocomputing 272, 294–305.

NCBI (1988). National Center for Biotechnology Information. National Library of Medicine, National Center for Biotechnology Information. Bethesda, Maryland. URL: https://www.ncbi.nlm.nih.gov/.

Neelakanta, G., Sultana, H., Fish, D., Anderson, J. F. and Fikrig, E. (2010). Anaplasma phagocytophilum induces Ixodes scapularis ticks to express an antifreeze glycoprotein gene that enhances their survival in the cold. J. Clin. Investig. 120, 3179–3190.

Palomares, M. L. D. and Pauly, D. [editors] (2020). SeaLifeBase. Vancouver, Canada. URL: https://www.sealifebase.ca/.

Patra, A. K., Chung, O., Yoo, J. Y., Kim, M. S., Yoon, M. G., Choi, J.-H. and Yang, Y. (2020). First draft genome for the sand-hopper Trinorchestia longiramus. Sci. Data, 7, 85.

Pratiwi, R., Malik, A. A., Schaduangrat, N., Prachayasittikul, V., Wikberg, J. E. S., Nantasenamat, C. and Shoombuatong, W. (2017). CryoProtect: a web server for classifying antifreeze proteins from nonantifreeze proteins. J. Chem. 2017, 9861752.

Pronk, P., Infante Ferreira, C. A. and Witkamp, G. J. (2005). A dynamic model of Ostwald ripening in ice suspensions. J. Cryst. Growth 275, e1355–e1361.

Prosdocimi, F., Linard, B., Pontarotti, P., Poch, O. and Thompson, J. D. (2012). Controversies in modern evolutionary biology: the imperative for error detection and quality control. BMC Genomics 13, 5.

R Core Team (2019). R: A language and environment for statistical computing. R Foundation for Statistical Computing, Vienna, Austria. URL: https://www.R-project.org/.

Raymond, J. A. (2014). The ice-binding proteins of a snow alga, Chloromonas brevispina: probable acquisition by horizontal gene transfer. Extremophiles 18, 987–994.

Raymond, J. A. and Remias, D. (2019). Ice-binding proteins in a chrysophycean snow alga: Acquisition of an essential gene by horizontal gene transfer. Front. Microbiol. 10, 2697.

Saier, B. (2002). Subtidal and intertidal mussel beds (Mytilus edulis L.) in the Wadden Sea: diversity differences of associated epifauna. Helgol. Mar. Res. 56, 44–50.

Sanmartín, I., Enghoff, H. and Ronquist, F. (2001). Patterns of animal dispersal, vicariance and diversification in the Holarctic. Biol. J. Linn. Soc. 73, 345–390.

Scholl, C. L., Tsuda, S., Graham, L. A. and Davies, P. L. (2021). Crystal waters on the nine polyproline type II helical bundle springtail antifreeze protein from Granisotoma rainieri match the ice lattice. FEBS J. Early View, 1–16.

Sevillya, G., Adato, O. and Snir, S. (2020). Detecting horizontal gene transfer: a probabilistic approach. BMC Genomics 21, 106.

Sorhannus, U. (2011). Evolution of antifreeze protein genes in the diatom genus Fragilariopsis: Evidence for horizontal gene transfer, gene duplication and episodic diversifying selection. Evol. Bioinform. 7, 279–289.

Storey, K. B. and Storey, J. M. (2013). Molecular biology of freezing tolerance. Compr. Physiol. 3, 1283–1308.

Sun, S., Ding, H., Wang, D. and Han, S. (2020). Identifying antifreeze proteins based on key evolutionary information. Front. Bioeng. Biotechnol. 8, 244.

Swanson, W. J. and Aquadro, C. F. (2002). Positive Darwinian selection promotes heterogeneity among members of the antifreeze protein multigene family. J. Mol. Evol. 54, 403–410.

Theede, H., Schneppenheim, R. and Béress, L. (1976). Antifreeze glycoproteins in Mytilus edulis? Mar. Biol. 36, 183–189.

Thompson, J. D., Higgins, D. G. and Gibson, T. J. (1994). CLUSTAL W: Improving the sensitivity of progressive multiple sequence alignment through sequence weighting, position-specific gap penalties and weight matrix choice. Nucleic Acids Res. 22, 4673–4680.

Tukey, J. W. (1949). Comparing individual means in the analysis of variance. Biometrics, 5, 99–114.

Usman, M. and Lee, J. A. (2019). AFP-CKSAAP: prediction of antifreeze proteins using composition of k-spaced amino acid pairs with deep neural network. In 2019 IEEE 19^th^ International Conference on Bioinformatics and Bioengineering (BIBE), pp. 38–43. IEEE.

Usman, M., Khan, S. and Lee, J. A. (2020). AFP-LSE: antifreeze proteins prediction using latent space encoding of composition of k-spaced amino acid pairs. Sci. Rep. 10, 7197.

Veltri, D., Malapi-Wight, M. and Crouch, J. A. (2016). SimpleSynteny: a web-based tool for visualization of microsynteny across multiple species. Nucleic Acids Res. 44, W41–W45.

Wang, S., Zhang, J., Jiao, W., Li, J., Xun, X., Sun, Y., Guo, X., Huan, P., Dong, B., Zhang, L. et al. (2017). Scallop genome provides insights into evolution of bilaterian karyotype and development. Nat. Ecol. Evol. 1, 0120.

Wang, S., Deng, L., Xia, X., Cao, Z. and Fei, Y. (2021). Predicting antifreeze proteins with weighted generalized dipeptide composition and multi-regression feature selection ensemble. BMC Bioinform. 22, 340.

Wang, Y., Graham, L. A., Han, Z., Eves, R., Gruneberg, A. K., Campbell, R. L., Zhang, H. and Davies, P. L. (2020). Carrot ‘antifreeze’ protein has an irregular ice-binding site that confers weak freezing point depression but strong inhibition of ice recrystallization. Biochem. J. 477, 2179–2192.

Warren, G. and Corotto, L. (1989). The consensus sequence of ice nucleation proteins from Erwinia herbicola, Pseudomonas fluorescens and Pseudomonas syringae. Gene, 85, 239–242.

Whelan, S. and Goldman, N. (2001). A general empirical model of protein evolution derived from multiple protein families using a maximum-likelihood approach. Mol. Biol. Evol. 18, 691–699.

Wiens, J. J. and Donoghue, M. J. (2004). Historical biogeography, ecology and species richness. Trends Ecol. Evol. 19, 639–644.

Wilbrandt, J., Misof, B., Panfilio, K. A. and Niehuis, O. (2019). Repertoire-wide gene structure analyses: a case study comparing automatically predicted and manually annotated gene models. BMC Genomics 20, 753.

Xiao, X., Hui, M. and Liu, Z. (2016). iAFP-Ense: an ensemble classifier for identifying antifreeze protein by incorporating grey model and PSSM into PseAAC. J. Membr. Biol. 249, 845–854.

Xu, H., Griffith, M., Patten, C. L. and Glick, B. R. (1998). Isolation and characterization of an antifreeze protein with ice nucleation activity from the plant growth promoting rhizobacterium Pseudomonas putida GR12-2. Can. J. Microbiol. 44, 64–73.

Yang, R., Zhang, C., Gao, R. and Zhang, L. (2015). An effective antifreeze protein predictor with ensemble classifiers and comprehensive sequence descriptors. Int. J. Mol. Sci. 16, 21191–21214.

Zhuang, X., Yang, C., Murphy, K. R. and Cheng, C.-H. C. (2019). Molecular mechanism and history of non-sense to sense evolution of antifreeze glycoprotein gene in northern gadids. PNAS, 116, 4400–4405.

